# Sub-district Costs and Efficiency of the *Re Mmogo Pholong* (“Together in Wellness”) Combination HIV/AIDS Prevention Intervention in the North West Province of South Africa

**DOI:** 10.1101/562926

**Authors:** Sebastian Kevany

## Abstract

**Background:** Re Mmogo Pholong (RMP) or “Together in Wellness”), was a combination prevention program to strengthen HIV prevention programming, community support mechanisms, community-based HIV testing, referral systems, and HIV prevention integration at the primary care level, thereby sustainably reducing HIV/AIDS transmission in the North West Province of South Africa. RMP included four overlapping components: situational analysis, community engagement and mobilization, community-based biomedical and behavioral prevention, and primary health care systems strengthening. In support of the PEPFAR country-ownership paradigm, we conducted costing analysis of the RMP combination HIV prevention program to determine data needed for potential transition of to local ownership.

**Methods:** We used standard costing methodology for this research.

**Results:** We found that cost per unit of output ranged from $63.93 (cost per person reached with individual or small group prevention interventions) to $4,344.88 (cost per health facility strengthened). The RMP intervention was primarily dependent on personnel costs. This was true regardless of the time period (Year 1 vs. Year 2) or activity (i.e. wellness days or events, primary health care strengthening, community engagement, and wellness clubs).

**Conclusions:** The development of labor-intensive rather than capital intensive interventions for low-income settings such as RMP was identified as being particularly important in treating and preventing HIV/AIDS and other health conditions in a sustainable manner. Costs were also observed to transition from international cost centers to in-country headquarters offices over time, in keeping with the transition of international to local responsibility required for sustainable PEPFAR initiatives. Such costing center evolution was also reflected by changes in the composition of the intervention, including (1) the redesign and re-deployment of service delivery sites according to local needs, uptake and implementation success and (2) the flexible and adaptable restructuring of intervention components in response to community needs.

## Background

### The Role of HIV Counseling and Testing and Combination Prevention

Combination HIV/AIDS prevention approaches, including multi-level initiatives that combine community mobilization, counseling and testing, and post-test support with other health services (e.g. Khumalo-Sakutukwa et al 2010) at the community-level have been shown to effectively increase utilization of HIV/AIDS and other health services in sub-Saharan Africa and elsewhere (Sweat et al 2011, Merson et al 2008). Economic costs and health-related quality of life outcomes of HIV treatment have also been examined in this regard and support the roll-out of these interventions (Sweat et al 2000, Creese et al 2002, Lalloo et al 2017). This is because, in part, comprehensive prevention integrates biomedical, behavioral, and structural strategies, with intervention components offered together increasing both (1) the likelihood of meeting the needs of diverse populations and (2) the potential for improved effectiveness due to synergies from complimentary approaches (Lippman et al 2015). Such innovative strategies to increase HIV/AIDS testing levels also reflect related HIV/AIDS counseling and testing (HCT) innovations in other sub-Saharan African countries (Maheswaran 2017), which have emerged as highly cost-effective programs.

Of the combination intervention components, HCT is the primary gateway into care and treatment, and is critical for stemming the spread of the epidemic (Khumalo-Sakutukwa et al 2011). Anchoring the provision of HCT services at the community level has, in turn, been found to increase testing uptake (Khumalo-Sakutukwa et al 2011) by reducing logistical, financial and social barriers such as lack of transportation, personal costs, and stigma. Expanding and integrating HCT activities with other behavioral and structural interventions has therefore been identified as an innovative approach to increasing overall prevention goals (Rotherham 2009), and has the potential to be more effective and cost-effective than HCT alone. In this regard, combined prevention and early treatment of other sexually transmitted infections (STIs) combined with HIV/AIDS efforts has for a number of years been a public health priority in South Africa, as cited in the National Strategic HIV/AIDS Plan (NSP) (South African Ministry of Health 2011).

### The Re Mmogo Pholong Combination HIV Prevention Program

*Re Mmogo Pholong* (RMP) was a program of the International Training and Education Center for Health (I-TECH), a collaboration between University of Washington (UW) and University of California, San Francisco (UCSF) in support of the South African Government to strengthen prevention programming, community support mechanisms, community-based HIV testing, referral systems, and HIV prevention integration at the primary care level. Such multi-component trials are increasingly considered to be the most effective and efficient means of addressing the HIV/AIDS epidemic in sub-Saharan Africa, reflecting their secondary capacities to improve, for example, drug adherence and retention in care (Granich et al 2009, Mangenah at al 2017). RMP also aimed to sustainably reduce HIV transmission in the Bojanala Platinum and Dr. Ruth Segomotsi Mompati Districts, North West Province, South Africa. RMP included four overlapping components: situational analysis, community engagement and mobilization, community-based biomedical and behavioral prevention (i.e. wellness days and wellness clubs), and primary health care systems strengthening. This comprehensive, multi-level and holistic strategy also aimed to sustainably reduce HIV/AIDS incidence and prevalence via integration with longer-term structural health care system change in the form of improved access to (and quality of) referral systems to higher-level care.

### Situational Analyses & Intervention Components

Situational analyses were conducted in both sub-districts in Year 1 (September 2011 to September 2012) (1) to understand the epidemic response in the community, (2) engage in community mapping, and (3) design programmatic interventions. Community mobilization and engagement took place through the development of community working groups (CWGs), engaging the South African Department of Health (DoH) for community entry and report-back on activities, and community mobilization strategies based on raising community awareness and engagement around HIV/AIDS prevention, and was considered an essential element of the broader program.

Related RMP service biomedical delivery was offered at “wellness days” and “wellness events”, building on evidence of their cost-effectiveness in the South African setting (Moodley et al, 2016). These provided HCT, referred and linked HIV-positive persons to care and treatment, provided pregnancy testing and family planning options, conducted rapid syphilis testing in pregnant women, provided syndromic management for STs and symptomatic screening and referral for TB to promote safer sexual behaviors, and emphasized the importance of consistent and correct condom use through health education. Wellness days were considered as extensions of public health care services, and referrals and linkages to Primary Health Centers (PHCs) were a primary focus.

Of note, those who received HCT also received support and guidance regarding personalized behavioral risk reduction, including partner and family testing, partner reduction, and disclosure of HIV and STI status. Individuals testing HIV-positive were referred to wellness clubs for psycho-social support, coping mechanisms, awareness of behavior change recommendations, and life skills development (e.g. patient goal setting, household and medical budget management, and involvement with food gardens and nutritional security). In each of the program components, linkage and referral to HIV care and treatment was a primary focus, and prevention amongst people living with HIV/AIDS (PLHIV) was an emphasis. We aimed to build on related research exploring the costs and outcomes of HIV/AIDS interventions in South African rural areas which often face very different operating environments and expenses compared to their urban counterparts (Mbonigaba and Oumar, 2017).

## Methods

### Costing the RMP Program

A key feature of the sustainability, transferability, and effectiveness of global health intervention implementation and roll-out involves the understanding and quantification of key costs (and related resources) required for service delivery. The costing process may also help to inform broader policy decisions related to resource allocation across HIV/AIDS treatment and prevention programmes (Marseille & Khan 2002), as well as providing a more detailed understanding of the key cost centers within program components. In the case of the RMP program, combined service delivery initiatives focused on provision of services via community engagement, wellness days, wellness clubs, and primary health center strengthening (the ‘cost centers’), across personnel, transport, facilities, supplies, and equipment cost categories.

The stratification of costs according to such categories is also related to geographical regions of expenditure (e.g. intervention sub-districts). We categorized and review related costs according to these classifications and approaches for the first two fiscal years of the intervention. Such “efficiency comparisons” represent a useful tool for local and international global health program managers and policymakers to determine returns on program investments (Jamison et al 2006). The use of output information from intervention monitoring and evaluation (M&E) activities, in conjunction with costing data, also presented opportunities to assess the efficiency of the implementation of RMP.

### Data Collection

Data for the costs incurred by the RMP program were obtained in several stages. UCSF analysts worked with I-TECH South Africa finance staff to extract South Africa-based expenditures from the local financial system for fiscal Year 1 (30^th^ September 2011 through 29^th^ September 2012) and Year 2 (30^th^ September 2012 through 29^th^ September 2013) in Excel format. Further consultation with the RMP management and program team at I-TECH South Africa (Seattle and in-country offices) helped to generate reports and code expenses related to RMP intervention development, implementation oversight, and other in-country costs. These data collection initiatives were complemented by ongoing communications with Seattle headquarters and in-country administrative and managerial staff in order to further inform categorization of intervention costs.

### RMP Site Visit

A costing site visit was conducted in February 2014 to observe service delivery at a two-day “wellness day” event in Bakubung Village, Moses Kotane sub-district, in the Bojanala Platinum District. This site visit allowed for (1) identification of any potential hidden costs not evident in the financial data and (2) a greater understanding of the operational needs of the intervention from a costing perspective.

### Data Entry and Stratification

All transactions for in-country cost data collection were originally presented in South African Rand (ZAR). Periodical average exchange rates were used to convert transaction costs to US dollars for the two fiscal years. Within each year, we then categorized costs by (1) location (e.g. Seattle headquarters office, in-country central office, and implementation sub-district); (2) intervention activity (e.g. headquarters oversight expenditures, in-country management and office costs, wellness days, wellness clubs, implementation science, primary health center strengthening, and community stakeholder engagement); and (3) economic resource categories such as capital (e.g. large, single purchase, equipment costs); personnel (e.g. salaries, benefits, and per diems,); utilities (e.g. electricity, communications); transport (e.g. vehicle maintenance); meetings (e.g. conference fees); oversight and support (e.g. maintenance and repair, or information technology and computing costs)^1^; space (e.g. office rental, conference facility rental); travel (e.g. air fare, shuttle service); medical supplies (e.g. HIV/AIDS testing kits); non-medical supplies (e.g. printing and copying or computer software); miscellaneous costs (e.g. freight and express, insurance, legal fees, and postage); and other costs (e.g. indirect costs related to administration).

### Inclusion of Output Data to Inform Efficiency Comparisons

As part of broader project oversight efforts, output data recorded during Year 2 was collected from project managers at the headquarters and field levels. Output information was then linked with site-specific costs aggregated by economic costing category (e.g. personnel) and divided by activity category (e.g. wellness days). These output values were then combined with component, economic category, and site-specific costs to generate a series of cost-efficiency ratios across sites. Shared costs or costs which could not be allocated to either sub-district were allocated proportionally by known site-specific costs. These results were, in turn, directly associated with (1) key costing groups (e.g. wellness days) and (2) geographical areas of service delivery (e.g. the Moses Kotane and Naledi sub-districts). Efforts to include the widest range of intervention outputs were made in keeping with current recommended practices (e.g. Padian et al 2012), helping to identify those sub-districts that performed with, for example, higher levels of productivity or lower costs.

Outputs related to key intervention activity categories of (1) community stakeholder engagement; (2) wellness days; (3) wellness clubs; and (4) primary health care facility strengthening. Outputs included, respectively, (1) number of communities with community working groups (CWGs); (2) number of wellness day events, the number of individuals reached with individual or small group HIV prevention interventions, the number of patients seeing a health care worker, the number of patients who received pre-test counseling, the number of HCT clients, and the number of individuals testing HIV-positive; (3) number of support groups completed; and (4) the number of efforts to build capacity for data collection, reporting, and analysis at the facility level.

### Data Analysis

Data were analyzed in Excel format in order to (1) compare trends in key cost centers between Years 1 and 2 and (2) to determine cost-efficiency in conjunction with output data. For costs, all line items from I-TECH South Africa accounting sources were classified according to time period, location, economic resource classification, and activity. First, in terms of time period, costs were classified as ‘Year 1’ (September 1^st^ 2011 to September 30^th^ 2012) and ‘Year 2’ (October 1^st^ 2012 to September 30^th^ 2013). Second, in terms of location, Year 1 and 2 costs were divided across the following sites: (1) University of Washington (Seattle) costs, (2) in-country headquarters costs, (3) Naledi sub-district-specific costs, and (4) Moses Kotane sub-district-specific costs.

Third, costs were divided by standard economic costing resource category (e.g. personnel, space, supplies, and travel costs). Fourthly, costs were allocated by activity component (e.g. wellness days and wellness clubs). Separately, for RMP outputs, information was transferred from standard reporting formats (e.g. Excel spreadsheets or tables embedded in Word and PDF documents) into a collective and inclusive Excel output database in which all outputs were quantified and divided into group (e.g. wellness clubs) and individual (e.g. number of HCT clients) level results. Efficiency comparisons were then completed by developing activity-specific sub-district-level costs and combining with key output categories.^2^ Finally, where multiple outputs were generated by a single activity category (e.g. the number of HCT clients and number of individuals reached for the wellness day activity), total component costs were applied to all output categories.

## Results

### Year 1 Costs

Year 1 represented the pre-implementation period of the intervention. At the site level, 49.9% of all Year 1 costs were attributable to University of Washington headquarters. University of Washington costs were driven primarily by personnel, oversight and support costs (52.0%). Year 1 costs were also divided across the following activities: (1) University of Washington headquarters support, (2) in-country management and office support, (3) wellness days, (4) wellness clubs, (5) implementation science, (6) primary health center strengthening, and (7) community stakeholder engagement.

Over sixteen percent (16.6%) of costs were attributable to in-country management and office support, while wellness days accounted for 14.6% of activity costs. In turn, in-country management costs were primarily generated by personnel costs (65.8%), while wellness day costs were primarily generated by equipment costs (53.2%). When stratified by economic category, key cost centers included personnel, oversight and support (64.1% of total costs) and equipment (9.3%). Of personnel costs, 47.1% of all costs were attributable to University of Washington costs, 17.0% to in-country office management, and 16.4% to implementation science. Of note, medical supplies accounted for only 1.2% of total intervention costs, non-medical supplies accounted for 0.7%, and space accounted for 4.2%.

### Year 2 Costs

At the site level, a total of 29.8% of Year 2 costs were attributable to the University of Washington headquarters, while a further 21.2% were attributable to the South Africa I-TECH in-country office. When considered by activity, Year 2 costs were primarily attributable to the University of Washington headquarters (29.8%), in-country management and office support (21.2%), wellness day (19.6%) and implementation science costs (16.7%).

Personnel, oversight and support costs, which made up 73.1% of all activity costs in Year 2, were focused on University of Washington headquarters costs (25.3%) and were almost equally attributable across wellness days (19.1%), implementation science (20.6%), and in-country management costs (19.8%). Finally, when stratified by economic resource category, costs were primarily generated by personnel, oversight and support (73.1%), travel (4.5%), and transport (2.6%).

### Output Results

Output results for Year 2 of the intervention were comparable between sub-districts for all group (e.g. wellness days, post-test ‘wellness’ clubs) and individual (e.g. number of HCT clients) level output categories (Table 1).

**Table 1:** Costs and Outputs of RMP by Activity.

| Activity / Cost Category | Community Stakeholder Engagement | Wellness Days |  |  |  |  |  | Post-Test Clubs | HSS |
| --- | --- | --- | --- | --- | --- | --- | --- | --- | --- |
| Output | Number of Communities Engaged | Number of Wellness Events | Individuals reached with individual or small group HIV prevention interventions | Patients Seeing Health Care Worker | Patients Pre-Test Counseled | Number of HIV Counselling and Testing (HCT) Clients | Patients HIV Positive | Number of Support Groups Completed | Number of primary health care facilities supported |
| Naledi | 10 | 100 | 3,025 | 2,865 | 1,619 | 1,474 | 76 | 18 | 4 |
| Moses Kotane | 22 | 121 | 4,142 | 4,137 | 3,614 | 2,771 | 152 | 29 | 9 |
| Cost by Activity (Naledi) | \$21,314.36 | \$264,802.20 | \$264,802.20 | \$264,802.20 | \$264,802.20 | \$264,802.20 | \$264,802.20 | \$65,469.07 | \$17,379.50 |
| Cost by Activity (Moses Kotane) | \$23,772.61 | \$268,216.95 | \$268,216.95 | \$268,216.95 | \$268,216.95 | \$268,216.95 | \$268,216.95 | \$76,203.65 | \$17,379.50 |
| Efficiency (Naledi) | \$2,131.44 | \$2,648.02 | \$87.54 | \$92.43 | \$163.56 | \$179.65 | \$3,484.24 | \$3,637.17 | \$4,344.88 |
| Efficiency (Moses Kotane) | \$968.83 | \$2,188.45 | \$63.93 | \$64.01 | \$73.27 | \$95.56 | \$1,742.12 | \$2,257.55 | \$1,931.06 |

Of note, the Moses Kotane sub-district recorded higher absolute levels of productivity for community engagement (number of CWG meetings), wellness days (number of HCT clients tested, number of wellness events, and number of individuals reached with small group HIV/AIDS prevention interventions), and wellness club activities (i.e. numbers of support groups completed). Associated output figures included 29 support groups completed at the Moses Kotane sub-district as compared to 18 at the Naledi sub-district, and 22 communities with CWGs as compared to 10. Naledi recorded 100 wellness events, from which 1,474 clients were tested, as compared to 121 wellness events at Moses Kotane at which 2,771 clients were tested. For wellness days, Moses Kotane sub-district was therefore associated with either comparable or significantly lower cost per wellness event, per HCT client, per individual reached, and per HIV-positive client diagnosed compared to associated costs in the Naledi sub-district.

### Sensitivity Analysis

Shared costs across Naledi and Moses Kotane per unit of output were determined on a proportional basis according to the distribution of assigned or “pre-allocated’ costs across communities. This approach was based on a proportional distribution of South African headquarters office, sub-district, and unknown costs. In order to ensure that our approach was appropriate and representative, we conducted a sensitivity analysis based on the equal distribution of headquarters, shared sub-district, and unknown costs across the two communities. Based on this analysis, costs per unit of output or outcome were not found to vary significantly. For example, cost per wellness event changed from $2,644.78 (Naledi) and $2,185.77 (Moses Kotane) to $2,648.02 and $2,188.45, respectively.

## Discussion

### Key Findings

Our key findings suggest that (1) the level of investment and responsibility for the RMP intervention may increase at the in-country level over time; (2) the cost per sub-district was highly variable and dependent on a range of environmental and structural factors such as operating environment and community buy-in and support; (3) personnel costs are a critical element of such interventions, and (4) the high level of indirect, management and oversight costs reflect the significant investment required at headquarters (in-country and international) versus field level.

These reflect and add to related findings such as those from related contemporary initiatives such as the SEARCH trial (Chamie et al 2012). Taken together, these findings may have important implications for PEPFAR expenditure planning, including (1) the need to explore additional investment for such interventions at the field level and (2) the desirability of labor-intensive interventions in resource-limited environments (Marseille and Kevany 2012). In regards the former, a number of community-level HIV interventions now focus their efforts and expenditure of field-level investments (Kevany et al 2013) and the possible opportunities for further investment in civil society, community support and stakeholder engagement at the sub-district level (Coates et al 2014) as strategies to ensure that programmatic responsibilities and investments continue to transition from the international to the local level under the country ownership paradigm (PEPFAR 2014).

### Changes in Costs over Time

Comparisons of Year 1 and 2 costs also reflect the transition from pre-implementation (with limited field activities) (Year 1) to a full implementation (Year 2) focus over time. For example, the University of Washington headquarters costs declined from 58.1% of total intervention costs to 29.8% between Year 1 and Year 2 of the intervention, reflecting increased in-country responsibilities during the full implementation phase. In the same way, many of those intervention costs related specifically to in-country implementation costs remained stable, or increased, from Year 1 to Year 2 of the intervention. These included wellness day costs (increasing from 14.6% to 18.0%) and primary health center strengthening costs (increasing from no investment in Year 1 to 19.6% of total intervention costs in Year 2).

Implementation science costs (e.g. situational analyses including impact evaluation pilot, and monitoring and evaluation activities) also increased between Year 1 and Year 2 from 12.5% to 16.7%, while costs of wellness clubs increased from 4.4% in Year 1 5.2% in Year 2. This overall shift in cost focus between donor and recipient country expenditure and investment (and from central office to field activity costs) is also reflected by changes in the proportion of combined in-country costs from 52.6% to 70.1% between Year 1 and Year 2, and is in keeping with the above-referenced country ownership paradigm (Kevany 2015).

### Cost Distribution across Sites

When reviewed by site, in both Year 1 and Year 2 of the RMP intervention costs were focused on the University of Washington -- with particular reference to personnel costs associated with program management -- indicating the importance of international involvement and support for the RMP intervention as a requirement for successful service delivery (with specific reference to the initial stages of the intervention) as well as protocol development and planning for the intervention, as reflected in related research findings (Coates et al 2014). Similarly, the primacy of South Africa headquarters management and office costs suggests that the RMP intervention was, during Year 1 and Year 2 of the intervention, dependent on high levels of centralized support and feedback, which may in turn have been related to the considerable logistical demands of a mobile health center based intervention, as also presented in prior cos-effectiveness analyses (Sweat el al 2000).

Of note, Seattle and in-country headquarters costs changed from $935,999.20 (58.17%) and $266,152.85 (16.53%) respectively during Year 1 of the intervention to $816,387.79 (29.83%) and $579,159.08 (21.16%) during Year 2 of the intervention – proportionately, an almost exact reversal of funding investments over time. This indicated a potentially successful transition of financial and organizational responsibility from the international to the in-country level, in keeping with other elements of the country ownership paradigm (Goosby 2014).

### Component and Activity-Related Costs

Analysis of activity-related costs suggested that the RMP intervention was highly dependent on the use of personnel. The development of labor-intensive rather than capital intensive interventions for low-income settings is particularly important in treating and preventing HIV/AIDS and other health conditions (Marseille & Kevany 2011), with particular reference to regions where a high availability of low-skilled personnel is available (as opposed to potentially scarce and expensive capital equipment). The associated importance of program management activities was also reflected in the personnel share of intervention costs. In addition, the low costs related to utilities and supplies also suggested that the RMP intervention would be easily transferrable to other low-income settings, a key feature of adaptable HIV/AIDS interventions (Kevany et al 2013).

### Cost per Unit of Output

Findings related to the cost per unit of output across each component of the RMP intervention may be of importance and use to policymakers in the South African and other contexts. For example, understanding and knowledge of the costs required to reach individuals or small groups with HIV/AIDS interventions (less than $100 per individual reached) might be compared with costs per HIV-positive identified ($3,484.24 in Naledi and $1,742.12 in Moses Kotane). Similarly, costs per patient accessing a health care worker were consistently less than $100 across communities and sensitivity analyses, while the cost per support group completed varied from $3,637.17 (Naledi) to $2,257.55 (Moses Kotane). Although such diverse elements of the intervention cannot be directly compared, and although each may have significantly different epidemic or health system strengthening impact, it may nonetheless be of interest to donors to identify those activities which produce, under the RMP approach, valuable outputs at low costs. Of note, these costs compared favorably to a number of recent studies on HCT-based interventions in South African and elsewhere (Menzies et al, 2009, Mangenah et al 2017).

### Explaining Differences in Efficiency across Sub-Districts

Program efficiency was found to be driven by productivity of outputs rather than by differences in sub-district-level costs. For example, the Moses Kotane sub-district recorded approximately double the level of outputs (compared to Naledi) across most indicators. A range of contextual, environmental and logistical considerations should be taken into account in this context. First, the Naledi sub-district was a geographically larger area, with populations spread over larger distances, which may affect uptake of services. By contrast, the Moses Kotane sub-district was more geographically concentrated, with less significant distances between population centers and RMP activities.

Second, the Moses Kotane sub-district was more accessible from the South Africa project headquarters office, meaning that travel expenses from Pretoria to Naledi (e.g. per diems and accommodation) were relevant to Naledi only. Third, the Moses Kotane sub-district was dominated by the mining industry, which impacted both population and migration. In this regard, several RMP activities took place at mining locations which resulted in higher numbers of participants and greater uptake in HCT per wellness day. Fourth, study staff at the field level observed that social, political and community level support for the RMP intervention diverged across communities, with greater community-level support for the intervention in the Moses Kotane sub-district. This was reflected in the number of communities engaged by the intervention (10 in the Naledi sub-district as compared to 22 in the Moses Kotane sub-district). Finally, the intervention was initially rolled out at the Moses Kotane sub-district and then moved to the Naledi sub-district two weeks later, which may have marginally contributed to differences in outputs across sub-districts.

### Policy Implications and Context

These findings are in keeping with key results from related literature. In comparable settings, combination HIV/AIDS interventions have been shown to focus resources on personnel and medical supplies, as well as transport and basic equipment, to facilitate field operations (Khumalo-Sakutukwa et al 2010, Marseille et al 2011). In addition, related results suggest that cost-efficiency declines with greater logistical demands, as well as in the context of recipient population and utilization considerations. In more rural areas, with more dispersed populations, these differences are often associated with additional transport and associated logistical costs (Zacariah et al 2006). However, given the lack of alternative resources in rural areas, combined with potentially high levels of detected and undetected HIV/AIDS prevalence, the case for continuing to provide services despite reduced cost-efficiency may be made on a health outcome, accessibility, and equity basis.

### Costing Perspective

The findings presented in this report represent a highly inclusive costing process, which encompassed all possible costs related to each component of the RMP intervention. Such holistic cost analyses, though still exclusive of patient costs, should be borne in mind when interpreting these results. Our approach is best represented by the health system perspective, which encompasses direct and indirect costs as well as costs borne by local headquarters offices and subcontracting support organizations. Other studies which present findings on the cost of, for example, HIV/AIDS counseling and testing from the provider perspective (with a focus on personnel time and medical and non-medical supplies) may therefore not be directly comparable in this regard (Ekwueme et al 2003).

### Limitations

There were a number of limitations to our analysis. Our review focused on a limited time period (Years 1 and 2 of the intervention). The possibility that Year 1 costs were not representative of usual operational costs due to start-up issues at this stage should therefore be considered. Our results also did not consider health outcomes beyond the number of HIV-positive patients identified through the wellness day process. The combination of cost and efficiency results with such results in future studies may help to further inform resource allocation and efficiency decisions.

## Conclusions

### Locally-Informed Resource Allocation Decisions

This analysis of the costs and outcomes of the RMP combination HIV/AIDS prevention program may help to inform associated resource allocation decisions and strategic planning at the in-country level. Our approach provided necessary data around activity-based costs, so that, in the country ownership context, the South African government could make necessary decisions related to efficiencies and effectiveness particularly in the context of changing proportions of funding investments between international and local management offices over time. Awareness of the key costs and efficiency findings of RMP may therefore have helped in-country actors such as the South African Department of Health to better anticipate associated resource requirements.

Findings related to costs across economic resource categories may have helped to sharpen the focus on the importance of personnel costs in the human resources for health (HRH) context (Schneider et al 2006), while our analysis of site-level costs may have helped to advance understanding of key cost centers -- by location, activity and economic category -- thereby helping to inform future implementers about the actual implementation costs of the RMP intervention. Finally, the inclusion of information about program outputs (e.g. number of wellness days held in each location or number of people receiving HCT), as related to the costs per intervention component (e.g. cost per client receiving HCT), may also have helped to inform interpretation of the cost and efficiency of such resource allocation decisions, ensuring that both funding agencies and recipient populations received optimal value for money from the public health perspective. Taken together, these results may have helped to ensure that PEPFAR-supported global health programs have continued to evolve towards more efficient design and implementation.

## Declarations

### Ethics approval and consent to participate

N/A

### Consent for publication

N/A

### Availability of data and material

N/A

### Authors’ contributions

SK is sole author

## Acknowledgements

N/A

## Footnotes

1 For the purposes of these analyses, personnel, oversight and support were grouped together as “personnel” and are referred to as “personnel” throughout this report.

2 Activity-specific costs for the Moses Kotane and Naledi sub-districts were generated via allocation of costs specifically assigned to each sub-district, as well as the proportional distribution of headquarters other cross-district costs.

